# mGluR5 PAMs rescue cortical and behavioural defects in a mouse model of CDKL5 deficiency disorder

**DOI:** 10.1101/2022.01.28.478021

**Authors:** Antonia Gurgone, Riccardo Pizzo, Alessandra Raspanti, Giuseppe Chiantia, Sunaina Devi, Noemi Morello, Federica Pilotto, Sara Gnavi, Leonardo Lupori, Raffaele Mazziotti, Giulia Sagona, Elena Putignano, Alessio Nocentini, Claudiu T. Supuran, Andrea Marcantoni, Tommaso Pizzorusso, Maurizio Giustetto

## Abstract

Cyclin-dependent kinase-like 5 (CDKL5) deficiency disorder (CDD) is a devastating rare neurodevelopmental disease without a cure, caused by mutations of the serine/threonine kinase CDKL5 highly expressed in the forebrain. CDD is characterized by early-onset seizures, severe intellectual disabilities, autistic-like traits, sensorimotor and cortical visual impairments (CVI). The lack of an effective therapeutic strategy for CDD urgently demands the identification of novel druggable targets potentially relevant for CDD pathophysiology.

To this aim, we studied metabotropic glutamate receptors 5 (mGluR5) for their important role in critical mechanisms involved in CDD, i.e.: synaptogenesis, dendritic spines formation/maturation and synaptic plasticity, and because mGluR5 function depends on the postsynaptic protein Homer1bc that is downregulated in the cerebral cortex of CDKL5^−/y^ mice. In this study, we reveal that CDKL5 loss tampers with (i) the strength of Homer1bc-mGluR5 binding, (ii) the synaptic localization of mGluR5 and (iii) the mGluR5-mediated enhancement of NMDA-induced neuronal responses. Importantly, we showed that the stimulation of mGluR5 activity by administering in mice specific positive-allosteric-modulators, i.e.: 3-Cyano-N-(1,3-diphenyl-1H-pyrazol-5-yl)benzamide (CDPPB) or RO6807794, corrected the synaptic, functional and behavioural defects shown by CDKL5^−/y^ mice. Notably, the cerebral cortex of 2 CDD patients show similar changes in the synaptic organization to mutant CDKL5 mice, including a reduced mGluR5 expression, suggesting that mGluR5 represent a promising therapeutic target for CDD patients.

## 1. Introduction

CDKL5 is a serine/threonine kinase highly expressed especially in the forebrain during the peak of synaptogenesis (Rusconi et al., 2008). CDKL5 phosphorylates several substrates and is involved in a broad variety of cellular processes such as gene expression, neuronal migration, axon outgrowth, dendritic morphogenesis, synapses development and function (Baltussen et al., 2018; Muñoz et al., 2018; Nawaz et al., 2016; Trazzi et al., 2016). In the nucleus CDKL5 has been shown to interact with epigenetic factors, such as methyl-CpG-binding protein 2 (MeCP2) and DNA Methyltransferase 1 (DNMT1) (Kameshita et al., 2008; Mari et al., 2005), nevertheless the role of CDKL5 in regulating gene expression is still not fully understood. Recently, several cytoplasmic targets of CDKL5 phosphorylation, including MAP1S, EB2 and ARHGEF2, have been identified pointing to a major role of this kinase in the control of cytoskeletal function. Moreover, CDKL5 has been found to accumulate at synapses where it can interact with the palmitoylated form of postsynaptic density protein-95 (PSD-95) (Zhu et al., 2013). The interaction with PSD95 facilitates the phosphorylation of the adhesion molecule netrin-G1 ligand (NGL-1) (Ricciardi et al., 2012) promoting the maturation of dendritic spines, i.e., the vast majority of glutamatergic postsynaptic sites in the forebrain, as well as the formation and function of excitatory connections. In addition, Barbiero et al. (2017) (Barbiero et al., 2017) showed that IQ motif containing GTPase activating protein 1 (IQGAP1) can interact with CDKL5 and thus mediate the formation of complexes with post-synaptic proteins such as PSD-95 or both AMPA- and NMDA-glutamatergic receptors. Interestingly, shRNA-mediated knockdown of CDKL5 can influence the synaptic expression of the GluA2 subunit (Tramarin et al., 2018) further highlighting that the involvement of CDKL5 in glutamatergic neurotransmission is yet to be unfolded.

To study the consequences of the lack of CDKL5 *in-vivo*, different CDKL5^−/y^ mouse lines have been recently generated (Amendola et al., 2014; Okuda et al., 2017; Wang et al., 2012). They all display a broad spectrum of behavioural abnormalities, including hind-limb clasping, motor hyperactivity, abnormal eye tracking, learning and memory deficits, and autistic-like phenotypes (Okuda et al., 2017), closely modelling human CDD (Demarest et al., 2019). Moreover, sensory defects such as tactile, visual and auditory impairments were recently revealed in CDD mouse models (Mazziotti et al., 2017; Pizzo et al., 2019; Wang et al., 2012). For example, cortical visual impairment (CVI), that is correlated with developmental delay in CDD patients (Demarest et al., 2019), is found in CDKL5 mutant mice starting from P27-P28 both in heterozygous and homozygous animals (Mazziotti et al., 2017; Wang et al., 2012).

Aberrant sensory processing in mice lacking CDKL5 is associated with severe abnormalities of the cerebral cortex, including altered dendritic arborization of pyramidal neurons, the downregulation of the postsynaptic scaffolding proteins PSD-95 and Homer, and the disruption of AKT-mTOR signaling (Amendola et al., 2014; Della Sala et al., 2016; Lupori et al., 2019; Pizzo et al., 2016; Wang et al., 2012). Moreover, we reported previously that CDKL5 plays a key role in the dynamic of dendritic spines turn-over in the primary somatosensory (S1) cortex (Della Sala et al., 2016) by promoting their stabilization. In addition, S1 cortex of CDKL5^−/y^ mice show impaired excitatory synaptic transmission and maintenance of long-term potentiation induced by theta-burst stimulation, emphasizing the role of CDKL5 in excitatory cortical connectivity (Della Sala et al., 2016; Pizzo et al., 2019).

Given all the above, we reasoned that by identifying druggable targets with relevant synaptic function is of pivotal importance to uncover novel therapeutic options for CDD. Here we report that both the expression and function of a member of group I metabotropic glutamate receptors, mGluR5, are abnormal in CDKL5^−/y^ mice cerebral cortex and that the administration of selective mGluR5 positive allosteric modulators (PAMs) can rescue synaptic, cellular, and behavioural defects shown by mutant mice.

## 2. Results

### 2.1 Altered mGluR5/Homer1bc organization in the cerebral cortex of Cdkl5^−/y^ mice

We focused the analyses on mGluR5 guided by mounting evidence pointing at their role in critical mechanisms involved in CDD such as synaptogenesis, dendritic spines formation/maturation and synaptic plasticity (Ballester-Rosado et al., 2016; Chen et al., 2012; Edfawy et al., 2019; Piers et al., 2012). mGluR5 needs to interact with Homer1bc to exert its signaling functions within the PSD (Giuffrida et al., 2005; Ronesi et al., 2012; Scheefhals and MacGillavry, 2018; Tu et al., 1999). Since Homer1bc is downregulated in the cortex of CDKL5^−/y^ mice (Pizzo et al., 2019, 2016), we evaluated the strength of mGluR5-Homer1bc binding in mutant mice. Intriguingly, co-immunoprecipitation (co-IP) assays of cortical synaptosomal fraction (fig.1A) revealed that the amount of mGluR5 immunoprecipitated with Homer1bc is significantly reduced in Cdkl5^−/y^ mice compared to Cdkl5^+/y^ animals (O.D mGluR5/Homer1bc * p < 0.05; fig 1B). We next assessed mGluR5 expression in the neuropil by performing immunofluorescence experiments on S1 cortices from Cdkl5^−/y^ and Cdkl5^+/y^ mice (fig. 1C). By using a fixation/staining protocol improved for postsynaptic protein localization (Morello et al., 2018; Pizzo et al., 2016), mGluR5 immunofluorescence (fig. 1C) resulted in discrete puncta that were found closely localized, but only rarely overlapping, with PSD-95^+^ puncta in agreement with previously reported perisynaptic localization of mGluR5 (Lujan et al., 1996). Interestingly, the quantitative analysis (fig. 1D) revealed that mGluR5-puncta density is strongly reduced in layers II-III and V of somatosensory (S1) cortex in Cdkl5^−/y^ mice compared to controls (layers II-III and layer V: Cdkl5^+/y^ vs Cdkl5^−/y^ * p < 0.05; fig. 1C-D). These findings indicate that the presence of CDKL5 is required for both mGluR5-Homer1bc binding and the synaptic localization of mGluR5.

**Figure 1.**
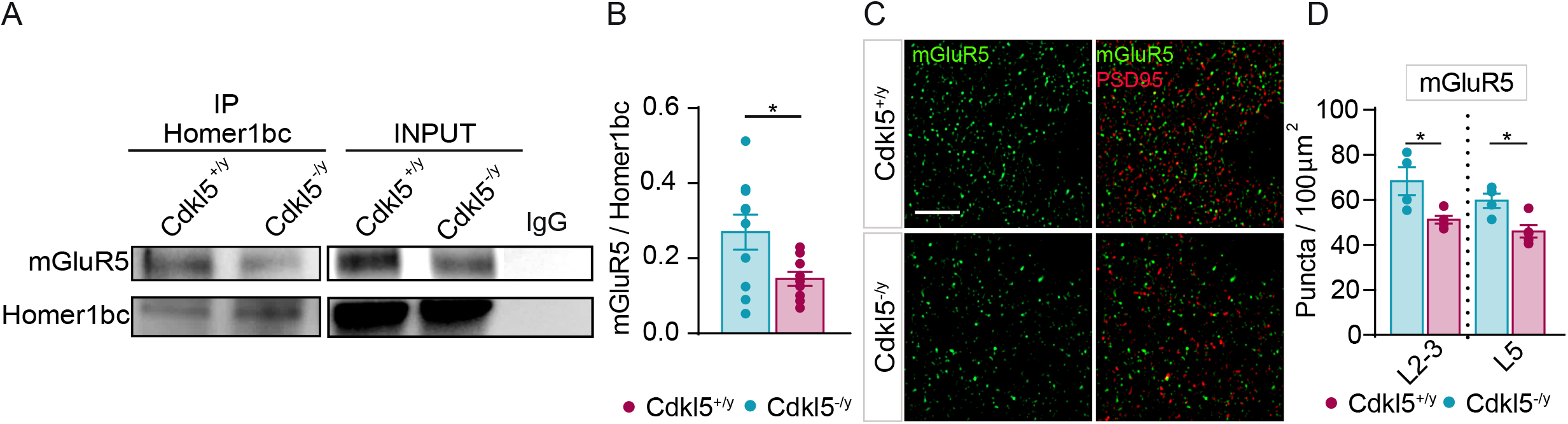
CDKL5 loss is responsible for both the disruption of mGluR5-Homer1bc interaction and the reduction of mGluR5 localization in the cortical neuropil. (**A**) Co-IP of cortical synaptosomal fraction (P2) from P56 mice by using anti-Homer1bc. IgG: control lane in the absence of antibodies. Immunoprecipitates and inputs were analyzed by immunoblotting for mGluR5 and Homer1bc. (**B**) Bar graphs showing Co-IP quantitation expressed as optical density (O.D.). (**C**) Confocal microscopy images showing mGluR5^+^ (green) and PSD-95^+^ (red) immunopuncta in layers II/III of S1 cortex (scale bar: 5 μm). (**D**) Bar graphs displaying the density of mGluR5^+^ puncta. Student T test *p < 0.05 (Co-IP: n = 8 IFL: n = 4).

### 2.2 mGluR5-mediated synaptic signaling is severely disrupted in Cdkl5^−/y^ cortical neurons

The reduced mGluR5-Homer1bc association that we found suggests that the receptor activity might be compromised (Aloisi et al., 2017; Kammermeier and Worley, 2007). To test this hypothesis, we started by recording spontaneous miniature excitatory postsynaptic currents (mEPSCs) in neuronal cultures of the S1 cortex from both Cdkl5^+/y^ and Cdkl5^−/y^ mice (fig. 2A-D, upper part), before and after mGluR5 activation. As we reported in a previous study (Della Sala et al., 2016), mEPSCs recorded from CDKL5 null neurons showed an increased inter-event interval (IEI) (Cdkl5^+/y^ vs Cdkl5^−/y^ * p < 0.05; fig. 2D) while the mean peak amplitude was similar between genotypes (Cdkl5^+/y^ vs Cdkl5^−/y^ p > 0.05; fig. 2C). Intriguingly, we found that when cortical neurons were stimulated for two minutes with the selective mGluR5 agonist DHPG (100 μm), mEPSCs IEI was significantly increased in Cdkl5^+/y^ cultures (fig. 2E, lower part; A) (Moult et al., 2006; Verpelli et al., 2011), but not in Cdkl5^−/y^ neurons (fig. 2E) indicating that CDKL5 loss impacts negatively on mGluR5 signaling in excitatory synaptic transmission. Next, we performed patch-clamp recordings in whole-cell configuration in neuronal cultures from S1 cortex and recorded NMDA currents elicited by NMDA (50 mM) alone (Marcantoni et al., 2020) or together with agonist DHPG (100 μm) as shown in Reiner et al., 2018 (Reiner and Levitz, 2018). Strikingly, Cdkl5^−/y^ cultures showed a significant reduction of INMDA with respect to Cdkl5^+/y^ neurons (Cdkl5^+/y^: 869.6 ± 85.4 pA, Cdkl5^−/y^: 544.2 ± 113.2 pA; ** p < 0.01; fig. 2F). Next, we found that DHPG increased INMDA in Cdkl5^+/y^ cells, (fig. 2G; see also Vicidomini et al., 2017) while, interestingly, it was not effective in Cdkl5^−/y^ neurons (fig. 2G), as illustrated by the sharp difference in the percentage of INMDA change between genotypes (Cdkl5^+/y^: 38.35%; Cdkl5^−/y^: –14.47%, p < 0.05. Fig. 2G). Strikingly, while 73% of Cdkl5^+/y^ cortical neurons (11/15 cells) showed potentiated INMDA after the application of DHPG, in most Cdkl5^−/y^ tested neurons we did not observe any effect of DHPG (10/14; 71%). These results disclose novel disrupted mechanisms of excitatory synaptic transmission caused by the absence of CDKL5, selectively involving metabotropic receptors signaling.

**Figure 2.**
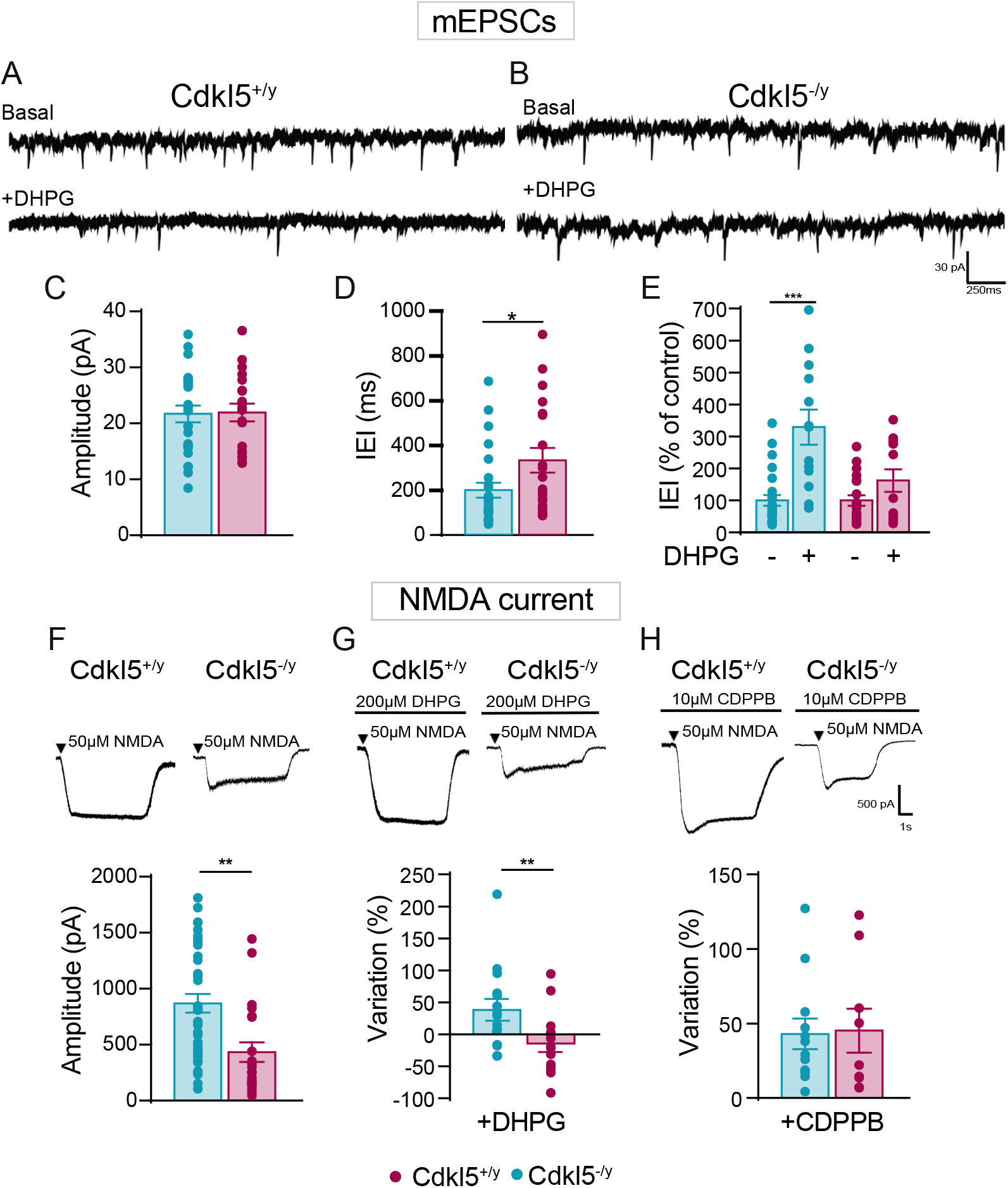
CDKL5 loss tampers with both mEPSCs and mGluR5-induced NMDA current. (**A**) Sample traces of miniature excitatory postsynaptic current (mEPSC) recorded from Cdkl5^+/y^ neurons (**A, upper part**) and Cdkl5^−/y^ neurons (**B, upper part**) and after the application of DHPG (**A, B lower part**). (**C-D**) Bar graphs showing the mean average amplitude (**C**) and the inter-event interval (IEI) of mEPSCs (**D**). (**E**) Bar graphs displaying the % change of IEI after the application of DHPG (100 μM). (**F**) Representative traces of currents obtained with patch-clamp recordings on S1 neurons cultures from Cdkl5^+/y^ and Cdkl5^−/y^ embryos after NMDA (50 μM) application (**upper part**), bar graphs showing differences of INMDA current between genotypes (**lower part**). (**G**) Representative traces of NMDA currents on S1 neurons after 2-min application of DHPG (100 μM-**upper part**); bar graphs showing the % change of INMDA after the application of DHPG (**lower part**). (**H**) Representative traces of NMDA after 2-min CDPPB + NMDA application (**upper part**), bar graphs showing the % change of INMDA current after the application of CDPPB (**lower part**). Student’s t-test, chi-square, two-way ANOVA followed by Fisher’s multiple comparison test, * p < 0.05, ** p < 0.01, *** p < 0.001 (mEPSC Cdkl5^+/y^ n = 22 cells, Cdkl5^−/y^ n = 28; minis+DHPG: n = 12 cells. NMDA: Cdkl5^+/y^ n = 36 cells, Cdkl5^−/y^ n = 23 cells; NMDA+DHPG Cdkl5^+/y^ n = 15 cells and NMDA+DHPG Cdkl5^−/y^ n = 14 cells; NMDA+CDPPB Cdkl5^+/y^ n = 12 cells; NMDA+CDPPB Cdkl5^−/y^ n = 9 cells).

### 2.3 CDPPB potentiates NMDAR current in cortical neurons lacking CDKL5

We and others have previously shown that in conditions where INMDA is not sensitive to DHPG, selective mGluR5 PAMs can instead elicit the strengthening of this current (Auerbach et al., 2011; Vicidomini et al., 2017). Among these, 3-Cyano-N-(1,3-diphenyl-1H-pyrazol-5-yl)benzamide (CDPPB) offers several advantages compared to agonist drugs such as higher subtype selectivity, reduced desensitization, and more subtle modulatory effects on receptor function (Chen et al., 2008). We first examined whether CDPPB produces an effect on cortical neurons lacking CDKL5 by measuring NMDA current. Intriguingly, we found that 2 min application of CDPPB (10 μM) preceding NMDA (50 μM) administration produced a comparable increase of INMDA (Fig. 2F, lower part) in both genotypes (Cdkl5^+/y^ 42.92%; Cdkl5^−/y^ 45.19%, fig. 2H) if compared to the average amplitude of INMDA measured after administration of NMDA alone. Interestingly, the majority of both Cdkl5^−/y^ and Cdkl5^+/y^ neurons showed potentiated INMDA after the application of CDPPB (Cdkl5^+/y^: 13/18, 78%; Cdkl5^−/y^ 10/12, 83%) resulting in a significantly higher percentage of Cdkl5^−/y^ neurons responding to CDPPB compared to DHPG treatment (chi-square Cdkl5^−/y^-DHPG: 29% vs Cdkl5^−/y^-CDPPB: 83% **** p < 0.0001). Thus, these results show that positive allosteric modulation can rescue mGluR5-dependent strengthening of NMDA-mediated activation in neurons lacking CDKL5.

### 2.4 CDPPB treatment ameliorates visual, sensorimotor and memory functions in Cdkl5^−/y^ mice

Encouraged by the positive effects that we obtained using CDPPB on synaptic currents, we evaluated the therapeutic potential of this PAM by treating mice with one intraperitoneal injection (i.p.) of CDPPB (3 mg/Kg), as in Vicidomini et al. (2017), and then exposing animals to a battery of tests.

We investigated cortical visual responses by transcranial intrinsic optical signal (IOS) imaging before and after CDPPB administration in the same animals. As expected from our previous data (Lupori et al., 2019; Mazziotti et al., 2017), baseline response amplitude of Cdkl5^−/y^ mice was strongly decreased compared to Cdkl5^+/y^ littermates (one way ANOVA ** p < 0.01; Tukey multiple comparison Cdkl5^+/y^ vs CDPPB-Cdkl5^−/y^ ** p < 0.01; Cdkl5^+/y^ vs vehicle-Cdkl5^−/y^ ** p < 0.01; vehicle-Cdkl5^−/y^ vs CDPPB-Cdkl5^−/y^ p = 0.92. Fig. 3A, B). After CDPPB treatment, visual responses approached Cdkl5^+/y^ levels (one way ANOVA ** p < 0.01; Tukey multiple comparison Cdkl5^+/y^ vs CDPPB-Cdkl5^−/y^ post injection p = 0.6; Cdkl5^+/y^ vs vehicle-Cdkl5^−/y^ post injection * p < 0.05) significantly increasing from their baseline values (two-way RM ANOVA; main effects not significant, interaction treatment*time* p < 0.05; Sidak multiple comparison: vehicle-Cdkl5^−/y^ post injection vs CDPPB-Cdkl5^−/y^ post injection * p < 0.05; CDPPB-Cdkl5^−/y^ baseline vs CDPPB-Cdkl5^−/y^ post injection * p < 0.05; vehicle-Cdkl5^−/y^ baseline vs vehicle-Cdkl5^−/y^ post injection p = 0.90). By contrast, visual response remained impaired in vehicle-treated mutant mice. These experiments indicate that the cortical visual impairment (CVI) shown by Cdkl5^−/y^ mice can be rescued by CDPPB treatment.

**Figure 3.**
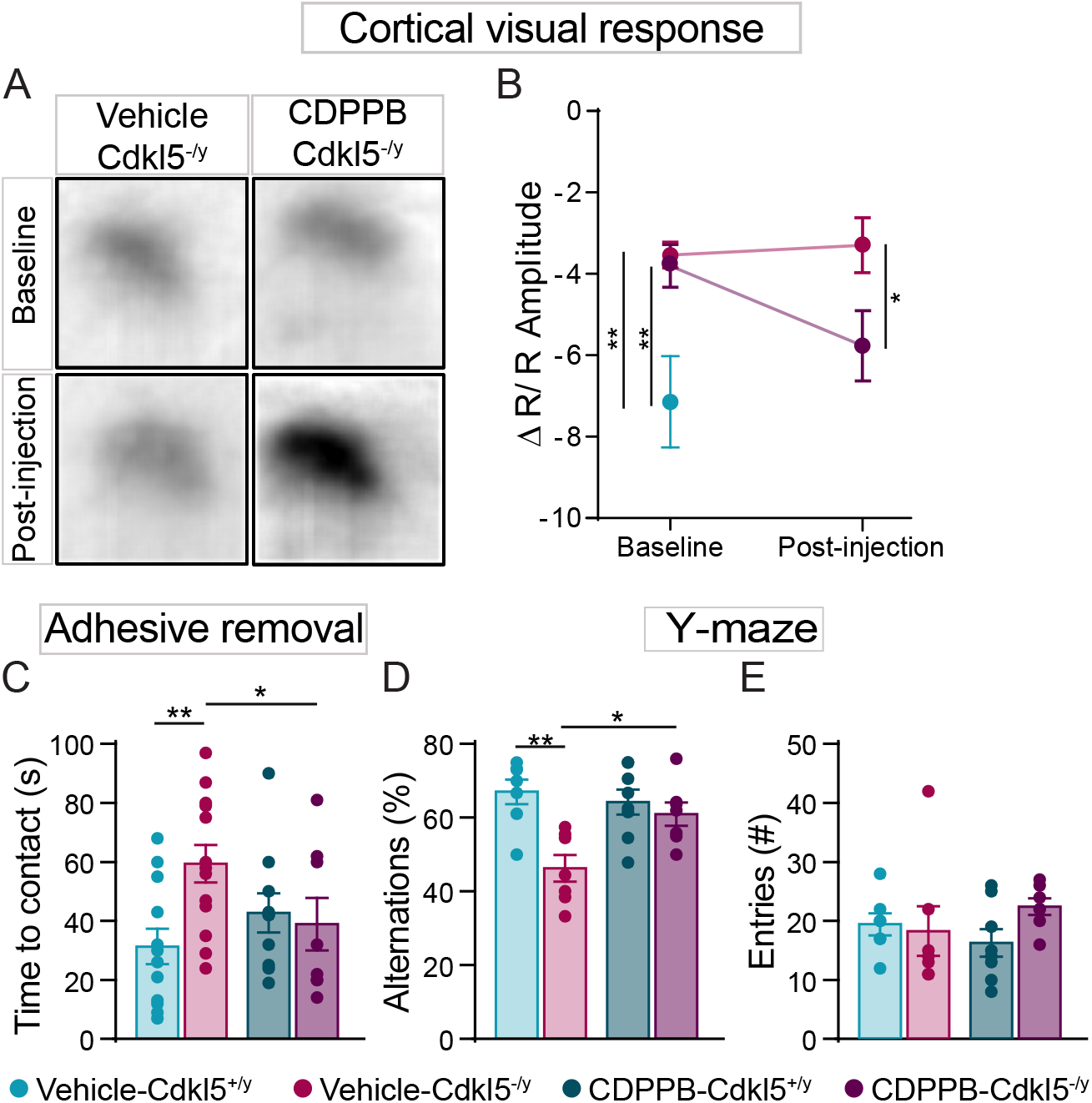
Acute CDPPB treatment rescues CVI, sensorimotor and memory deficits in Cdkl5^−/y^ mice. (**A**) Samples images showing differences of IOS evoked responses in vehicle- and CDPPB-treated Cdkl5^−/y^ mice. (**B**) Trajectory of the IOS amplitude in vehicle-Cdkl5^+/y^, vehicle-Cdkl5^+/y^ and CDPPB-Cdkl5^−/y^ treated mice. (**C**) Bar graphs showing contact latency with the tape placed under mice’s forepaw. (**D, E**) Bar graphs showing the percentage of the correct alternations (**D**) and the number of entries (**E**) made by Cdkl5^+/y^ and Cdkl5^−/y^ mice, treated with either vehicle or CDPPB, in the Y-maze. Two-way ANOVA followed by Sidak or Bonferroni’s multiple comparison test, * p< 0.05, ** p< 0.01 (IOS: vehicle-Cdkl5^+/y^ n = 3, vehicle-Cdkl5^−/y^ n = 8, CDPPB-Cdkl5^−/y^ n = 6; behavioural tests: vehicle-Cdkl5^+/y^ n = 12, vehicle-Cdkl5^−/y^ n = 13, CDPPB-Cdkl5^+/y^ n = 8, CDPPB-Cdkl5^−/y^ n = 7).

When we assessed sensorimotor responses by using the adhesive tape-removal test (Bouet et al., 2009; Komotar et al., 2007), we found that Cdkl5^−/y^ mice display a significant increase in the time-to-contact the tape compared to Cdkl5^+/y^ mice (vehicle-Cdkl5^+/y^ vs vehicle-Cdkl5^−/y^ ** p < 0.01; fig. 3C). Importantly, a single CDPPB injection produced a reduction of the latency exclusively in mutant mice whose performance became similar to controls (vehicle-Cdkl5^+/y^ vs CDPPB-Cdkl5^−/y^ p > 0.4; fig. 3C). Finally, the effect of CDPPB was assessed in the Y-maze paradigm for working memory, a feature known to be impaired in Cdkl5^−/y^ mutant mice (Fuchs et al., 2014). First, we found that the number of the correct spontaneous alternations is decreased in Cdkl5^−/y^ mice compared to Cdkl5^+/y^ animals (vehicle-Cdkl5^+/y^ vs vehicle-Cdkl5^−/y^ ** p < 0.01; fig. 3D), confirming previous observations. Intriguingly, CDPPB rescued working memory defects in Cdkl5^−/y^ mice by normalizing the frequency of spontaneous alternations (vehicle-Cdkl5^+/y^ vs CDPPB-Cdkl5^−/y^ p > 0.4; fig. 3D), without altering the performance of Cdkl5^+/y^ mice. No difference in the total number of arms entries were found between genotypes under either treated or untreated conditions (fig. 3E). Thus, these data indicate that the action of CDPPB can reverse atypical functional responses, such as CVI and sensorimotor defects, as well as memory impairment shown by Cdkl5^−/y^ mice.

### 2.5 mGluR5 PAMs rescue both synaptic and activity defects in Cdkl5^−/y^ cerebral cortex

In parallel with the observed behavioural and functional rescues, we found that the acute CDPPB treatment produced a normalization of the number and organization of postsynaptic sites as well as of the activity in primary cortices of Cdkl5^−/y^ mice. CDPPB increased the density of Homer1bc^+^ puncta in both S1 and V1 cortices of Cdkl5^−/y^ mice (S1: layers II-III and layer V vehicle-Cdkl5^−/y^ vs CDPPB-Cdkl5^−/y^ ** p < 0.01. V1: layers II-III and layer V vehicle-Cdkl5^−/y^ vs CDPPB-Cdkl5^−/y^ * p < 0.05; fig. 4A,B), reproducing Cdkl5^+/y^ mice conditions (S1 and V1: layer II-III and layer V: vehicle-Cdkl5^+/y^ vs CDPPB-Cdkl5^−/y^ p > 0.3; fig. 4A,B). Intriguingly, CDPPB treatment also normalized mGluR5^+^ puncta density in both S1 and V1 cortices of Cdkl5^−/y^ mice (S1: layers II-III and layer V: vehicle-Cdkl5^−/y^ vs CDPPB-Cdkl5^−/y^ *** p < 0.001; vehicle-Cdkl5^+/y^ vs CDPPB-Cdkl5^−/y^ p > 0.3. V1: layers II-III and layer V: vehicle-Cdkl5^−/y^ vs CDPPB-Cdkl5^−/y^ * p < 0.05. S1 and V1: vehicle-Cdkl5^+/y^ vs CDPPB-Cdkl5^−/y^ p > 0.3; fig. 4C,D). Finally, the density of cells expressing ARC, an immediate-early gene induced by mGluR5 activation (Ménard and Quirion, 2012; Wang and Zhuo, 2012), was restored in the V1 cortex of CDKL5-mutants after a single CDPPB administration (layers I-VI: vehicle-Cdkl5^+/y^ vs vehicle-Cdkl5^−/y^ ** p < 0.01; vehicle-Cdkl5^−/y^ vs CDPPB Cdkl5^−/y^ *** p < 0.001; fig. 4E,F).

**Figure 4.**
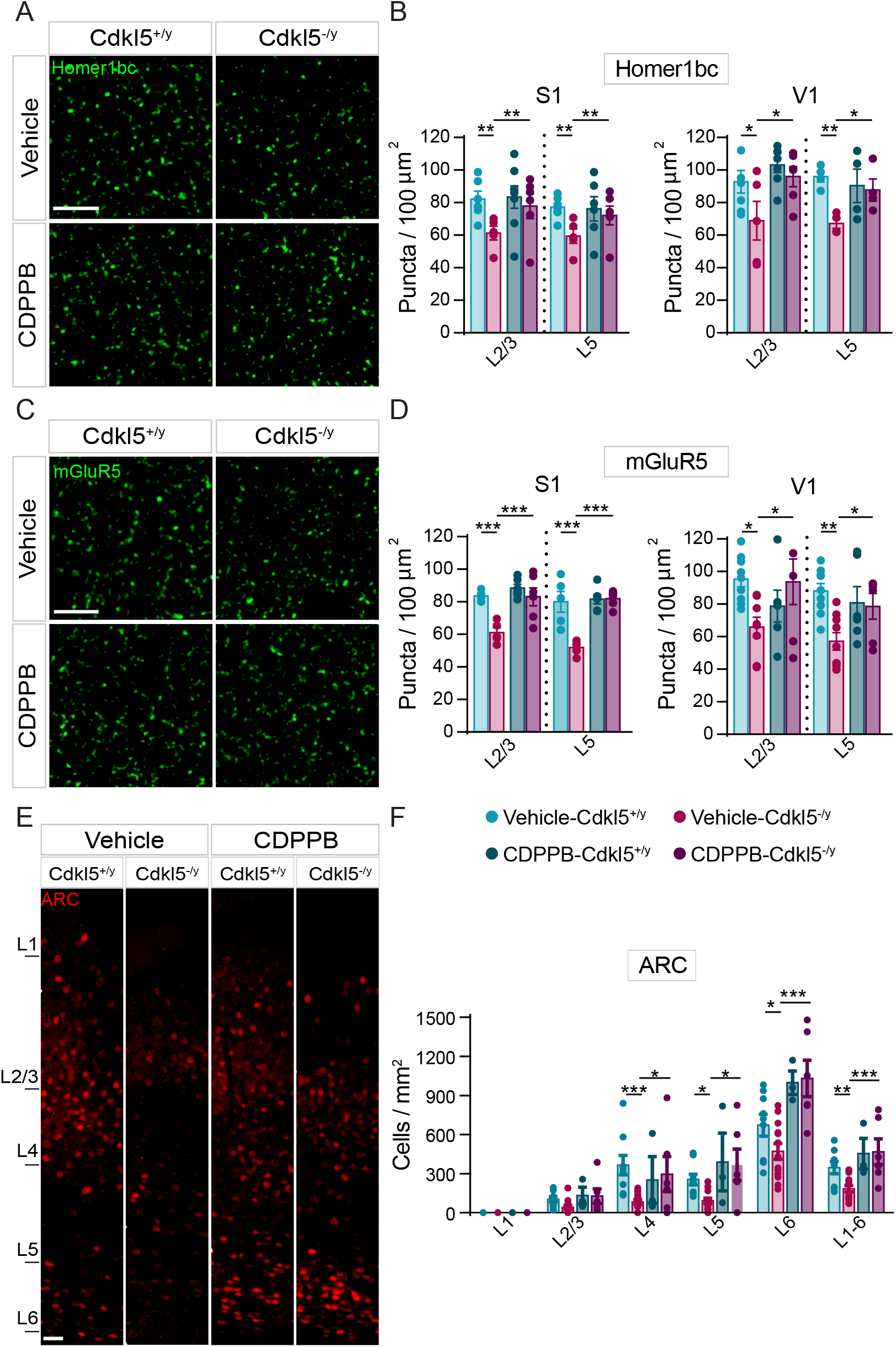
Structural defects exhibited by Cdkl5^−/y^ mice cortices are rescued by an acute CDPPB injection. (**A, C**) Representative confocal images showing Homer1bc^+^ and mGluR5^+^ puncta in layer II-III of S1 cortex from either vehicle- or CDPPB-treated mice (scale bar: 5 μm). (**B, D**) Bar graphs showing both Homer1bc^+^ (**B**) and mGluR5^+^ (**D**) immunopuncta density in layers II-III and V of both S1 and V1 cortices in either vehicle- or CDPPB-treated mice. (**E**) Confocal images of ARC immunostaining on coronal sections of the V1 cortex from mice treated with vehicle or CDPPB (scale bar: 25 μm), and relative ARC^+^ cells density quantitation (**F**) throughout the cortical layers. Two-way ANOVA followed by Fisher’s multiple comparison test, *p < 0.05, ** p < 0.01, *** p < 0.001; (n = 6 animals for each genotype).

In order to increase the reproducibility of our study, we treated another group of Cdkl5^−/y^ and Cdkl5^+/y^ animals with a different mGluR5 PAM, the RO6807794 (RO68) compound (Kelly et al., 2018). Two hours after an i.p. injection with RO68 (0.3 mg/kg as in Kelly et al., 2018), we found that the density of Homer1bc^+^ puncta in S1 cortex of Cdkl5^−/y^ mice was increased (S1: layers II-III and layer V vehicle-Cdkl5^−/y^ vs CDPPB-Cdkl5^−/y^ * p < 0.05. V1: layers II-III and layer V vehicle-Cdkl5^−/y^ vs CDPPB-Cdkl5^−/y^ * p < 0.05; fig. S1 A-B) to reproduce Cdkl5^+/y^ mice conditions (S1 layer II-III and layer V: vehicle-Cdkl5^+/y^ vs CDPPB-Cdkl5^−/y^ p > 0.3; fig. S1 A-B). Intriguingly, RO68 was also able to restore neuronal activity in S1 cortex in Cdkl5^−/y^ mice (Fig. S1 C) throughout cortical layers (vehicle-Cdkl5^+/y^ vs vehicle-Cdkl5^−/y^ *** p < 0.001; vehicle-Cdkl5^−/y^ vs RO68-Cdkl5^−/y^ *** p < 0.001), as indicated by the c-Fos^+^ cell density count (Fig. S1 D; see also Pizzo et al., 2016), that reached the magnitude of Cdkl5^+/y^ mice (vehicle Cdkl5^+/y^ vs CDPPB-Cdkl5^−/y^ p > 0.05). These results strongly support the idea that the atypical organization, both structural and molecular, of the neural circuits in the cerebral cortex of Cdkl5^−/y^ mutants can be rescued by activating mGluR5-mediated signaling.

### 2.6 The effects of a protracted CDPPB treatment in Cdkl5^−/y^ mice are long-lasting

To assess the therapeutic potential of mGluR5 activation, we treated animals for five consecutive days with CDPPB and 24 hours after the last injection animals were behaviourally tested and then sacrificed for brain analyses. We found that, after this protracted treatment, the density of Homer1bc^+^ puncta remained increased in both upper and deeper layers of the S1 cortex in mutant mice (layers II-III and layer V: vehicle-Cdkl5^−/y^ vs CDPPB-Cdkl5^−/y^ p < 0.01; fig. 5A, B), and that its value was no longer different from control (layers II-III and layer V: vehicle-Cdkl5^+/y^ vs CDPPB-Cdkl5^−/y^ p > 0.3; fig. 6B). In contrast, no effect of CDPPB on Homer1bc expression was found in Cdkl5^+/y^ animals (layers II-III and layer V: vehicle-Cdkl5^+/y^ vs CDPPB-Cdkl5^+/y^ p > 0.9; fig. 5A, B). Next, we analysed hind-limb clasping, a sign displayed shown by Cdkl5^−/y^ mice (Amendola et al., 2014; Terzic et al., 2021; Trazzi et al., 2018). In line with previous studies, vehicle-treated mutants showed increased hind-limb clasping compared to Cdkl5^+/y^ littermates (vehicle-Cdkl5^+/y^ vs vehicle-Cdkl5^−/y^ p < 0.001; fig. 5C -Amendola et al., 2014) whereas after CDPPB treatment Cdkl5^−/y^ mice spent significantly less time clasping their hind paws (vehicle-Cdkl5^−/y^ vs CDPPB-Cdkl5^−/y^ p < 0.01; fig. 5C), although CDPPB did not produce a complete normalization (vehicle-Cdkl5^+/y^ vs. CDPPB-Cdkl5^−/y^ p < 0.01; fig. 5C). Finally, we evaluated the effects of sub-chronic CDPPB treatment on cortical activity by analysing c-Fos expression. The effect of CDPPB treatment was subtle (vehicle-Cdkl5^+/y^ vs vehicle-Cdkl5^−/y^ p < 0.01; vehicle-Cdkl5^−/y^ vs CDPPB-Cdkl5^−/y^ p = 0.09; fig. 5 D,E) but sufficient to normalize c-Fos levels in mutant mice (vehicle Cdkl5^+/y^ vs CDPPB-Cdkl5^−/y^ p > 0.09; fig. 5E). Taken together, these data indicate that protracted CDPPB treatment is accompanied by long-lasting positive effects in Cdkl5^−/y^ mice, a result consistent with the design of a therapeutic protocol for CDD targeting mGluR5.

**Figure 5.**
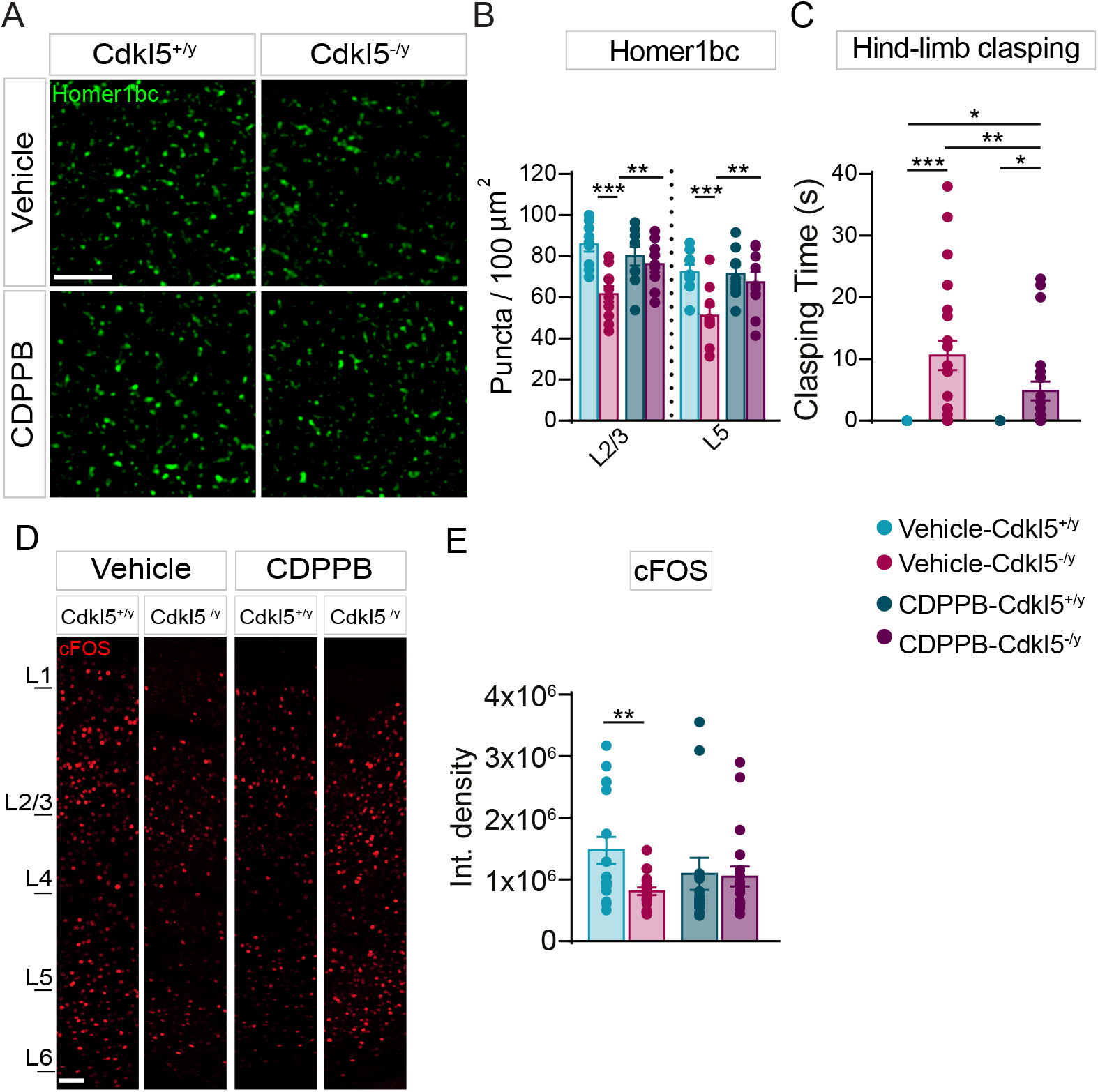
The subchronic treatment with CDPPB produces lasting effects in Cdkl5^−/y^ mice. (**A**) Representative confocal images of Homer1bc^+^ (red) immunofluorescence in layers II-III of the S1 cortex. (**B**) Bar graphs showing Homer1bc^+^ puncta density in layers II-III and V of the S1 cortex of either vehicle- and CDPPB-treated mice. (**C**) Bar graphs showing time spent clasping in vehicle- and CDPPB-treated mice. (**D**) Representative images of c-Fos immunoreactive cells in S1 of vehicle- and CDPPB-treated mice (scale bar 50 μm). (**E**) Bar graphs showing the integrated intensity analysis of c-Fos immunofluorescence in the S1 of vehicle- or CDPPB-treated mice. Two-way ANOVA followed by Fisher’s LSD: * p < 0.05, ** p < 0.01, *** p < 0.001 (Homer1bc^+^ puncta: n = 9 for each group; clasping and c-Fos vehicle-Cdkl5^+/y^ n = 34, vehicle-Cdkl5^−/y^ n = 23, CDPPB-Cdkl5^+/y^ n = 17, CDPPB-Cdkl5^−/y^ n = 23).

**Figure 6.**
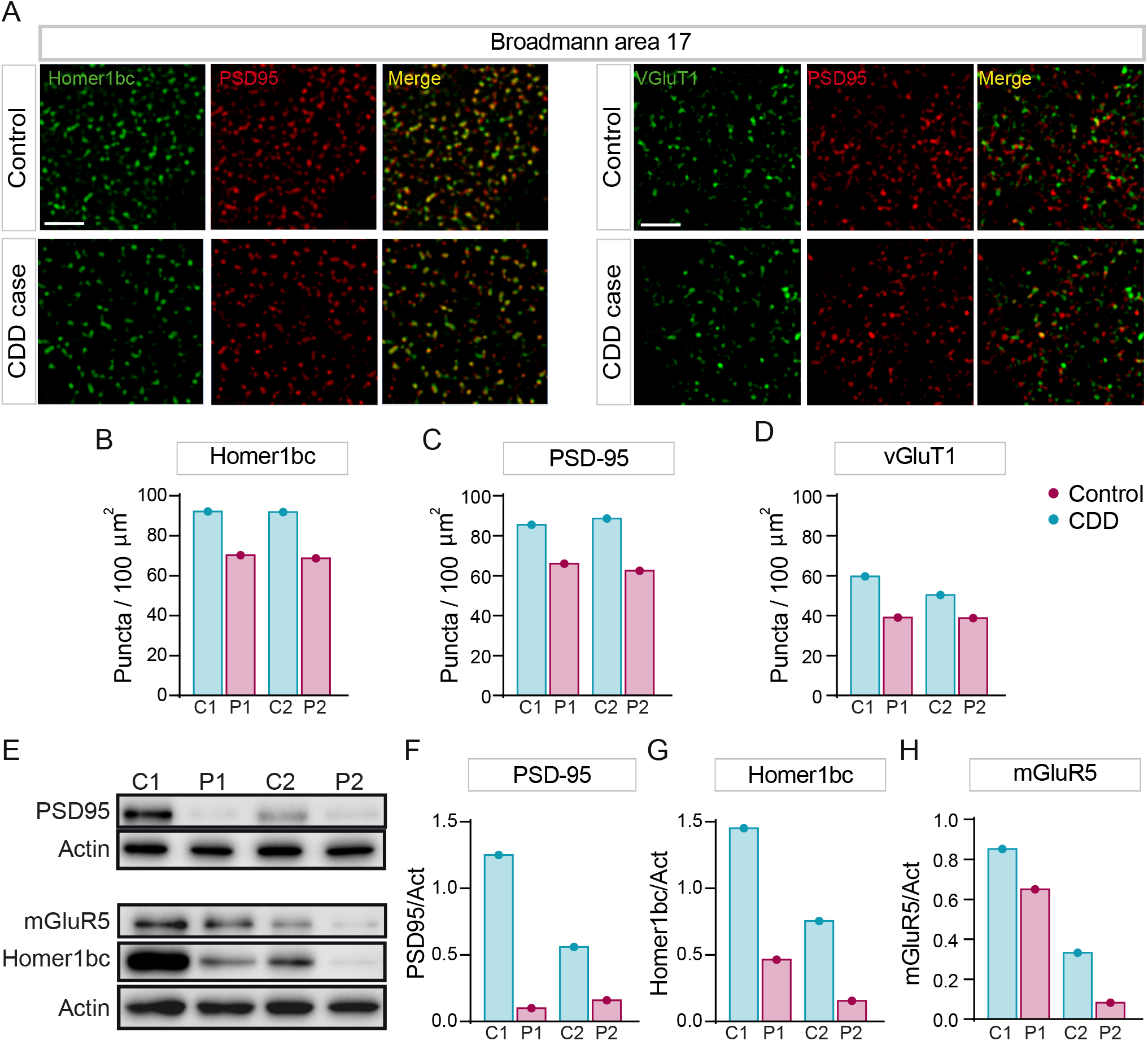
Aberrant expression of excitatory synaptic proteins in the BA17 cortex of CDD patients. (**A)** Illustrative confocal images taken from layers II-III of the BA17 cortex. (**A**) PSD-95^+^(red), Homer1bc^+^(green), VGluT1^+^(green) immunofluorescence puncta. Note the virtually complete overlapping of PSD-95 and Homer1bc immunofluorescence (scale bar: 5 μm). (**B, C, D**) Bar graphs showing the analysis of puncta density in layers II-III of BA17 cortices. (**E**) Western blotting showing the expression of PSD95, Homer1bc and mGluR5 in lysates from BA17 cortices. (**F-H**) Bar graphs displaying the optical density (O.D.) analysis of PSD95 (**F**), Homer1bc (**G**) and mGluR5 (**H**) expression. Student’s t-test, * p < 0.05, ** p < 0.01 (C1 = F, 4 years old; P1 = F, 5.7 years old; C2 = F, 29 years old; P2 = F, 30 years old).

### 2.7 The BA17 cortex of CDD patients recapitulates the mGlur5 defects shown by Cdkl5^−/y^ mice

Finally, to assess the translational potential of our findings, we examined excitatory synaptic structures in the 2 postmortem CDD patient brains available worldwide that we obtained from the Harvard Brain Tissue Resource Center (Belmont; USA). These experiments were performed on sections from the primary visual cortex (BA17) of CDD cases and age/sex-matched neurotypical subjects (NTs). Intriguingly, the results showed a clear reduction of both postsynaptic proteins PSD-95^+^ and Homer1bc^+^ as well as of the presynaptic marker VGluT1^+^, irrespective of case age (5 and 30 years old), compared to NTs (fig. 6A)). Moreover, although a statistical analysis was not performed with only 2 cases, the quantification of the immunopuncta revealed a reduction in the cortices of CDD patients compared to NTs (fig. 6A-D), indicative of an overall reduction of glutamatergic synapses. We next evaluated Homer1bc, PDS-95 and mGluR5 expression by western blotting on BA17 cortex lysates. Intriguingly, as shown in figures 6 E-H, the BA17 area from CDD samples showed a robust reduction of their expression compared to NTs. Although derived from a limited dataset, these results suggest for the first time that both structural and molecular signatures of CDKL5 loss largely overlap between mice and human cortical connectivity and support the translational potential of a mGluR5-directed therapeutic strategy.

## 3. Discussion

It is urgent to find therapeutic targets that shall be rapidly translated into treatments for CDD, a devastating condition without corrective options. In this study, we focus our attention on mGluR5, a group I metabotropic glutamate receptor highly expressed in the cerebral cortex of both mice and humans (Ferraguti & Shigemoto, 2006). To properly function, mGluR5 requires binding with Homer1bc (Aloisi et al., 2017), a scaffolding protein that is severely downregulated in the cerebral cortex of both CDKL5^−/y^ mice and CDD patients (Pizzo et al., 2019, 2016; fig. 6A) as well as in iPSCs-derived neurons from CDD patients (Negraes et al., 2021).

We here show for the first time that CDKL5 plays a role in the expression of mGluR5 in the cerebral cortex of both CDD patients and CDKL5 mutant mice, an effect likely produced by the defective formation of mGluR5-Homer1bc complexes at synapses as indicated by our data. Moreover, we find that synaptic transmission, both basal and NMDA-mediated, is altered in S1 neurons lacking CDKL5 and that it is unresponsive to the modulation normally produced by the selective mGluR5 agonist DHPG. Because Shank1, by forming complexes with Homer1bc, PSD-95 and NMDAR, promotes the cooperation between NMDAR and mGluR5 signaling machineries (Ango et al., 2000; Hering and Sheng, 2001; Tu et al., 1999), our electrophysiological evidences strongly suggest that lack of CDKL5 tampers with the synergistic cooperation between these glutamatergic receptors. This effect is likely produced by a reduced amount of Homer1bc recruited in the postsynaptic density in the absence of CDKL5 which, in turn, results in an atypical postsynaptic localization/stabilization of mGluR5. Interestingly, aberrant NMDA receptors signaling have been previously reported by Okuda and colleagues (2017) in the hippocampus of a different CDKL5 mutant mice line showing severe NMDA-dependent epileptic seizures due to the incorrect postsynaptic accumulation of GluN2B-containing NMDA receptors (Okuda et al., 2017). Similar results have been obtained in the hippocampus of the Cdkl5^R59X^ knock-in CDD mouse model (Yennawar et al., 2019). Altogether, although with some differences, these findings further support the idea that CDKL5 plays a crucial role in the correct localization/function of glutamate receptors, both ionotropic and metabotropic, at the synapse. Remarkably, an aberrant expression and function of mGluR5 has been reported in several neurodevelopmental diseases such as Fragile X, Phelan McDermid syndrome, Tuberous sclerosis (TSC) and Rett syndrome (Aloisi et al., 2017; Auerbach et al., 2011; Gogliotti et al., 2016; Vicidomini et al., 2017) further supporting the primary role of mGluR5 signaling as common deranged pathway in monogenetic forms of neurodevelopmental disorders.

The reduced expression/function of mGluR5, combined with relevant synaptic and behavioural signs shown by CDKL5 mutant mice, provided us with solid bases for attempting the first preclinical assessment of mGluR5 PAMs efficiency for treating CDD that we report in this study. Intriguingly, our results revealed that an acute treatment with CDPPB is effective in restoring several endophenotypes and behavioural signs produced by CDKL5 loss both *in-vitro* and *in-vivo*. Our data show that in primary cortical neuronal cultures, CDPPB can restore mGluR5-mediated potentiation of NMDA currents in Cdkl5^−/y^ pyramidal neurons. Considering the negative response of NMDA current to DHPG treatment that we report in mutant neurons, the effect of CDPPB is surprising and still without a clear pharmacological explanation, although it closely replicates what has been found previously in Shank3-KO neurons (Vicidomini et al., 2017). Furthermore, the present findings strongly suggest that CDPPB treatment can facilitate the functional maturation of dendritic spines in the absence of CDKL5, as it increases the synaptic expression of both Homer1bc and mGluR5, two crucial molecular determinants of spine formation and stabilization (Oh et al., 2013; Sala et al., 2003). These synaptic effects are reflected by beneficial outcomes at the functional and behavioural level in mutant animals that are mostly relevant in the context of CDD. Our data, showing for the first time that both CVI and overall cortical activity can be rescued by CDPPB treatment in Cdkl5^−/y^ mice, strengthens the translational value of our preclinical results. As a matter of fact, recent data indicate that CVI is correlated with reduced milestone achievement in CDD patients and therefore CVI can be used in the clinic as a solid biomarker for CDD diagnosis, progression, and treatment with biunivocal translational validity(Demarest et al., 2019; Mazziotti et al., 2017).

Interestingly, our findings indicate that mGluR5 signaling greatly suffers from the lack of CDKL5, but it does not become completely non-functional. In support of this idea, our data show that the protracted treatment with CDPPB in CDKL5-null mice produces long-lasting rescuing effect on both the density of dendritic spine-like structures and cortical c-Fos expression in the cerebral cortex as well as on the hindlimb-clasping phenotype. Thus, although further studies are needed to dissect out the mechanisms of CDPPB action on excitatory synapse signaling, our results encourage further testing of mGluR5 PAMs in animals modelling CDD and offer hope for a future use of these compounds in the clinic. Our set of data obtained with another mGluR5 PAM, the RO68 compound, further strengthen this idea. Considerably, RO68 has the clinically relevant advantage that it can be dissolved in salina with an extremely low percentage of detergent (i.e.: Tween-80) and has a higher potency compared to other mGluR5 PAMs. Remarkably, RO68 is efficacious even at very low concentrations (i.e.: 0.3 mg/kg), thus reducing the risk of toxicity, as we show in this study where this compound was able to rescue neuroanatomical, functional and behavioural signs of CDKL5 mutant mice, and as it was previously shown in a mouse model of TSC (Kelly et al., 2018).

Finally, the positive action of mGluR5 PAMs on the molecular organization of postsynaptic structures is encouraging in view of the data we have obtained from the only two post-mortem CDD brains available. Remarkably, we show, for the first time, that the lack of CDKL5 induces the disorganization of the excitatory synaptic compartment in the BA17 cortical area of CDD human brains both the localization and expression of several synaptic molecules (i.e. VGluT1, Homer1bc, PSD-95 and mGluR5) seems negatively affected, as we and others previously reported in CDKL5-null mice (Pizzo et al., 2019, 2016; Trazzi et al 2018; Amendola et al., 2016). These results, when confirmed on a larger group CDD brains, shall contribute to disclose the connectivity impairments of the primary visual cortex underlying CVI in these patients (Demarest et al., 2019) and strengthen the face-validity of CDKL5^−/y^ mice in closely modelling the neuropathological signs of CDD. Importantly, our data indicate that the synaptic abnormalities and mGluR5 downregulation occurring in human CDD patients are potentially rescuable by positive allosteric modulation of mGluR5.

In further support of our findings, Negraes et al. have found similar synaptic defects in iPSCs-derived cortical neurons from CDD patients (Negraes et al., 2021). In apparent contrast from our observation, they found an increased mGlurR5-PanHomer association in CDD human organoids (Negraes et al., 2021) while we revealed that mGluR5-Homer1bc binding, an association crucial for this receptor function, is decreased in CDKL5 mutant mice. The most parsimonious explanation of this discrepancy arises primarily either the different technical approaches or the experimental models used (i.e., mice brain vs CDD human organoids). Moreover, in Negraes et al. no discrimination between different Homer isoforms has been attempted, although it is known that the binding between mGluR5 and Homer1bc or Homer1a produces opposite effects on mGluR5 membrane expression and function (Menard and Quirion, 2012; Shiraishi-Yamaguchi and Furuichi, 2007; Bertaso et al., 2010). Hence, the enhanced mGlur5-Pan-Homer interaction could be produce by an increased association with Homer1a, thus not ruling out a decreased of mGluR5-Homer1bc binding as revealed by our study.

In conclusion, we believe that our findings on the efficacy of mGluR5 activation pave the way for including these receptors as a promising therapeutic target for CDD. Our results also suggest that an early-onset and prolonged regime of mGluR5 activation has the potential to stably revert the morphofunctional defects shown by adult CDKL5 mutants. Finally, this study further supports previous indications that abnormalities of mGluR5 signaling represents a convergent pathway for multiple neurodevelopmental diseases, a solid hallmark now including CDD.

## 4. Materials and Methods

### Animals and pharmacological treatment

Animal care and handling throughout the experimental procedures were conducted in accordance with European Community Council Directive 2010/63/UE for care and use of experimental animals with protocols approved by the Italian Minister for Scientific Research (Authorization number 38/2020-PR) and the Bioethics Committee of the University of Torino, Italy. Animal suffering was minimized, as was the number of animals used. Mice for testing were produced by crossing Cdkl5^−/x^ females with Cdkl5^−/y^ males or with Cdkl5^+/y^ males. Littermate controls were used for all the experiments. After weaning, mice were housed 4 per cage on a 12 h light/dark cycle (lights on at 7:00 h) in a temperature-controlled environment (21 ± 2°C) with food and water provided ad libitum. For this study, 8-weeks old (post-natal day 56) Cdkl5^−/y^ and Cdkl5^+/y^ males were used. Because we did not observe any noticeable interindividual phenotypic or metabolic (e.g. weight and health condition scores) difference among the mouse cohorts used in this study, no inclusion/exclusion criteria were adopted besides age (PND56) and sex (male) of the animals. In pharmacological rescue experiments, animals were treated with the selective positive allosteric modulator (PAM) of mGluR5 3-cyano-N-(1,3-diphenyl-1H-pyrazol-5-yl)benzamide (CDPPB; Tocris, UK) that was diluted in saline solution containing 5-10% final concentration of DMSO and polyethylene glycol 400 (DMSO: PEG 400 = 1:9) 5,23. Acutely treated mice received an intraperitoneal injection (ip) of either CDPPB (3 mg/kg) or vehicle 5 at 9.00 am and then put back in their home cage for 1 hour before being behaviourally tested. For sub-chronic administration, animals were treated for five consecutive days and, 24 hours after the last injection, tested and after one hour from the behavioural test, the animals were sacrificed. All mice were subsequently sacrificed for brain analyses. All analyses presented in this study were carried out by investigators who were blinded to the animal’s/neuronal genotype or treatments.

### Synaptosomal fraction preparation

Adult mice (PND 56) were killed by decapitation, the entire cortex was rapidly removed and tissue was processed as in 61,62. The tissues were homogenized in ice-cold lysis buffer (0.32 M sucrose, and HEPES 1X at pH 7.4 and 1 mM EGTA, 1mM Na-Orthovanadate, 1 mM DTT, phenylmethylsulphonyl fluoride and 1 mM sodium fluoride and protease inhibitors (SIGMAFAST™ Protease Inhibitor Cocktail Tablets, EDTA-Free), using a glass Teflon tissue grinder. The homogenates were centrifuged at 1000 g for 10 min at 4°C. After discarding the nuclear pellet, the supernatant was centrifuged at 12,500 g for 20 min at 4°C. The P1 fraction was then washed with the same initial volume of lysis buffer and underwent further spin (20 min; 12,500 g). The pellet obtained was the crude cortical synaptosomal fraction (P2) that was resuspended in 400 μl of RIPA buffer (NP-40 1%, Na deoxycholate 0,25%, EDTA 0.5M, NaCl 5M, Tris pH8 1M, SDS 10%) and stored at −80°C. Protein concentration of the synaptosomal fraction was determined with the Bio-Rad protein assay kit.

### Co-immunoprecipitation assay

50 μg of proteins from the crude synaptosomal (P2) fractions were incubated for 1 hour at 4 °C in RIA buffer (200 mM NaCl, 10 mM EDTA, 10 mM Na2HPO4, 0.5% Nonidet P-40 supplemented with 0.1% SDS) and protein A/G-agarose beads (Santa Cruz, Dallas, TX, USA) as pre-cleaning procedure. The beads were then let to sediment at the bottom of the tube and the supernatant was collected. Primary antibody (Homer1bc) was added to the supernatant before leaving to incubate overnight (O/N) at 4 °C on a rotating wheel, then protein A/G-agarose beads were added and incubation continued for 2h at RT. Beads were then collected by gravity and washed three times with RIA buffer before adding sample buffer for SDS–polyacrylamide gel electrophoresis (SDS–PAGE) and heating the mix al 95°C for 10 min. Beads were pelleted by centrifugation an supernatants were separated using 4–15% SDS–PAGE precast gels (Biorad, Italy) (Mellone et al., 2015).

### Human postmortem brain tissue

The CDKL5 specimens were provided by the Harvard Brain Tissue Resource Center, Belmont (USA). Case P1 was a 5.7-year-old female with a frameshift mutation (c.2153_2154dupTG) in exon 15 of CDKL5 gene that results in a premature stop codon. Case P2 was a 30-year-old female with a deletion of exons 1-3 in CDKL5 gene. Control samples were obtained from the University of Maryland, Baltimore. Case C1 was a 4 years old female whereas case C2 was a 29 years old female. The sections analysed are from the BA17 occipital region of human cerebral cortex. Some sections were lysate for western blot analyses, others for immunofluorescences. While, six 30 μm-thick sections were pull down for each samples, and processed with RIPA buffer (NP-40 1%, Na deoxycholate 0,25%, EDTA 0.5M, NaCl 5M, Tris pH8 1M, SDS 10%).

### Western blotting

Lysates both immunoprecipitates and the inputs (50% of the total P2 lysates) and from human brains were boiled in SDS sample buffer, separated by SDS–PAGE and the proteins were then blotted to PVDF membrane following a standard protocol (Grasso et al., 2017). Next, PVDF membranes were blocked in BSA 5% for 1h and incubated with the primary antibodies (see table 2) O/N at 4°C. After washes with TBS 0.1% Tween 20, the membranes were incubated with the appropriate secondary antibodies (anti-mouse or anti-rabbit, 1:5000; Sigma, Italy) for 1h at RT. The chemiluminescent signal was visualized using Clarity™ Western ECL Blotting Substrates (Bio-Rad; Italy) and acquired with Bio-Rad ChemiDocTM Imagers (Bio-Rad; Italy) and analysed with Image J software (NIH, Usa).

**Table 1.**
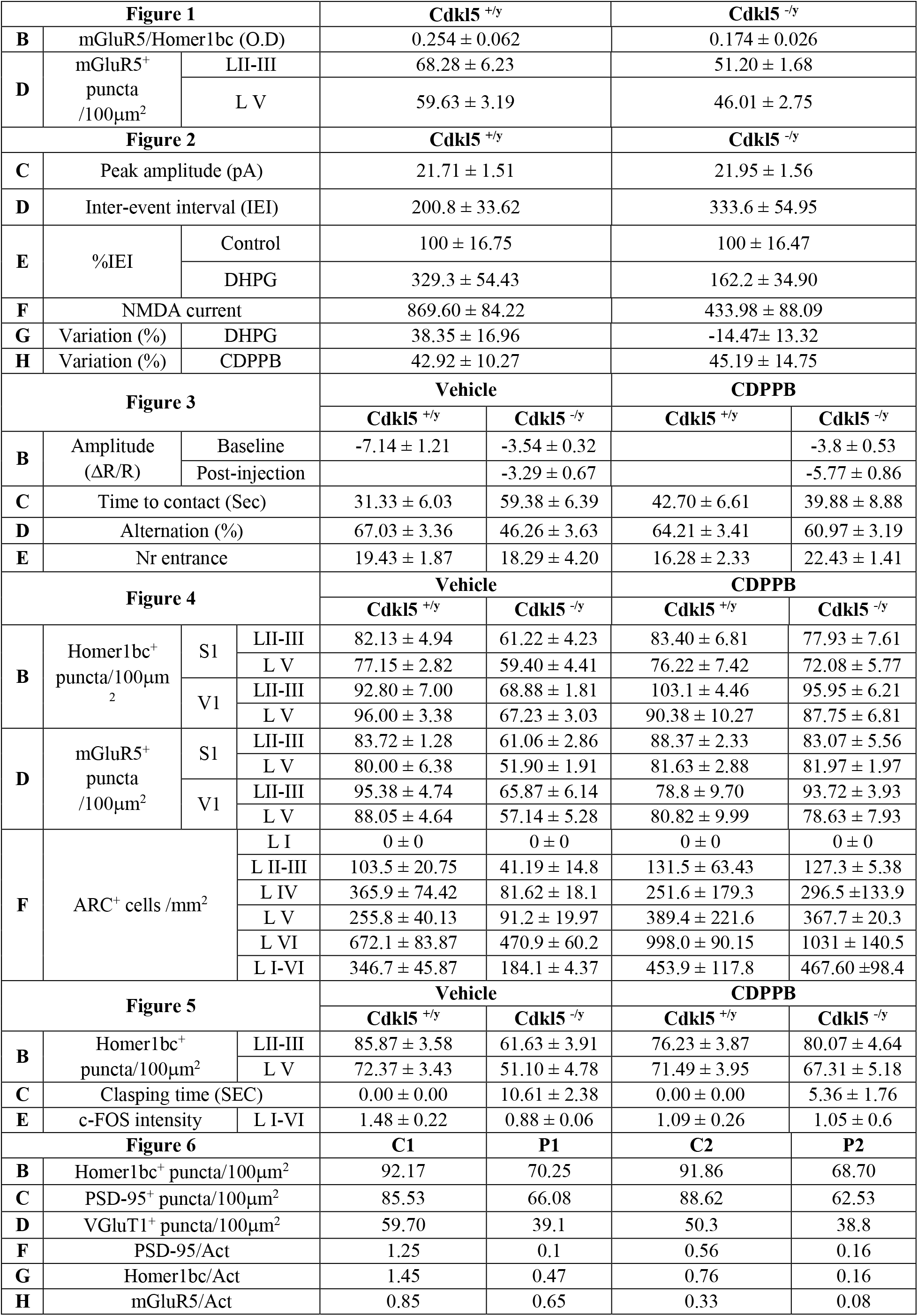
Mean ± SEM values for each statistical analysis

**Table 2.** List of antibodies used

| Primary Antibody | Species of origin | Working dilution |  |  | Supplier and catalog no. |
| --- | --- | --- | --- | --- | --- |
|  |  | WB | IF | IP |  |
| Homer1bc | Rabbit | 1:1000 | 1:500 |  | Synaptic System, Germany, cod. 160 023 |
| Homer1bc | Mouse |  |  | 1 mg | Santa Cruz Biotechnology, cod. sc-25271 |
| mGluR5 | Rabbit | 1:500 | 1:250 |  | Millipore, Germany, USA cod. AB5675 |
| ARC | Rabbit |  | 1:500 |  | Synaptic System, Germany, cod. 156 003 |
| c-FOS | Rabbit |  | 1:1500 |  | Santa Cruz Biotechnology, USA, cat. sc-52 |
| PSD-95 | Mouse | 1:500 | 1:250 |  | Neuromab; CA, USA, Clone K28/43 |
| VGluT1 | Guinea pig |  | 1:5000 |  | Millipore, Germany, cat. 5905 |

### Immunofluorescence procedures

<u>Cerebral cortical tissue</u>: for synaptic proteins detection, mice were anesthetized using a mix of tiletamine/zolazepam (40mg/kg) and xilazine (4-5 mg/kg) and then decapitated 20. The brains were rapidly excised and manually cut in coronal slabs that were fixed by immersion in ice-cold paraformaldehyde (4% in 0.1M phosphate buffer, PB, pH 7.4) for 30 min. After fixation, tissue slabs were rinsed in PB 0.1M, cryoprotected by immersion in sucrose-PB 0.1M solutions (10, 20 and 30%), cut in 20-μm sections with a cryostat, mounted on gelatine-coated slides and stored at −20°C until immunolabeling was performed as in 20. For c-Fos, ARC and Homer1bc immunodetection after CDPPB subchronic treatment, animals were anesthetized tiletamine/zolazepam (40mg/kg) and xilazine (4-5 mg/kg) and transcardially perfused with about 10 ml of 0.1M PBS followed by 80 ml of ice-cold 4% paraformaldehyde in 0.1M PB. After the brains were dissected, they were kept in the same fixative solution O/N at 4°C, cryoprotected by immersion in raising sucrose-PB 0.1M solutions (10, 20 and 30%), cut into 30 μm sections with a cryostat and stored at −20°C in a cryoprotectant solution containing 30% ethylene glycol and 25% glycerol until use. Cryosections were subsequently processed free-floating by immersion in 0.1M PBS solution containing 3% normal donkey serum (NDS) and 0.5% Triton X for 1h followed by an O/N incubation at 4°C with the primary antibodies (see table 2). The following day the sections were rinsed with 0.1M PBS and incubated with the appropriate fluorescent secondary antibodies (anti-mouse or anti-rabbit 1:1000; Jackson ImmunoResearch, West Grove, PA, USA) for 1h at RT. The sections were washed three times with PBS, mounted on gelatine-coated glass slides and cover slipped with Dako fluorescence mounting medium (Dako Italia, Italy).

<u>Human postmortem brain tissue</u>: immunofluorescence was performed on flash-frozen sections. Serial sections (20 μm) were cut by using a cryostat, mounted on superfrost slides and stored at −80°C until immunolabeling was performed. Before starting the immunofluorescence, sections were fixed in cold methanol for 1 min. The sections were then processed for double immunofluorescence by using in combination anti-Homer1bc and anti-PSD-95 primary antibodies and the appropriate fluorescent secondary antibodies (anti-mouse or anti-rabbit 1:1000; Jackson ImmunoResearch, West Grove, PA, USA) following the same protocol used for mouse brain tissue (see Pizzo et al., 2016).

### Images acquisition and analysis

For brain sections analyses, the layers of the mouse primary somatosensory and visual (S1 and V1) cortices were identified as previously reported (Tomassy et al., 2014; van Brussel et al., 2009). Synaptic immunofluorescence puncta, for both mouse and human brain tissue, were analysed on 5 serial optical sections (0.5 μm Z-step size) acquired from layers 2-3 and 5 of S1 and V1 with a laser scanning confocal microscope (LSM5 Pascal; Zeiss, DE) using a 100× oil objective (1.4 numerical aperture) and the pinhole set at 1 Airy unit. The density of the immunopositive puncta was determined by manual count followed by density analysis (puncta/100 μm^2^) with Imaris (Bitplane, Zurich, CH) and Image J (USA) softwares. Synaptic puncta were included if present in at least two consecutive optical sections.

For ARC and c-Fos immunofluorescence analyses, confocal images of the mouse S1 and V1 cortices were acquired in at least three corresponding coronal brain sections from at least six animals per group with a 20× objective using a 1-μm Z-step. Digital boxes spanning from the pial surface to the corpus callosum were superimposed at matched locations on each coronal section of V1 and divided into 10 equally sized sampling areas (bins; layer I: bin 1; layer II/III: bins 2–3; layer IV: bins 4–5; layer V: bins 6–7; layer VI: bins 8–10) as in (Tomassy et al., 2014). The density of ARC^+^ cells was determined by manual count in each bin using Image J software and expressed as cells/mm2, while intensity values of c-Fos^+^ cells were obtained using a dedicated Image J tool (integrative density) to analyse Z-stack projected images (Sum value).

### Chronic IOS Imaging

<u>Surgery</u>: for chronic IOS preparations, adult mice were anesthetized and maintained with isoflurane (respectively 3 and 1%), placed on a stereotaxic frame and head fixed using ear bars. Body temperature was controlled using a heating pad and a rectal probe to maintain the animals’ body at 37°C. Local anaesthesia was provided using subcutaneous lidocaine (2%) injection and eyes were protected with dexamethasone-based ointment (Tobradex, Alcon Novartis). The scalp was removed, and the skull carefully cleaned with saline. Skin was secured to the skull using cyanoacrylate. Then a thin layer of cyanoacrylate is poured over the exposed skull to attach a custom-made metal ring (9 mm internal diameter) centred over the binocular visual cortex. When the glue dried off, a drop of transparent nail polish was spread over the area to ameliorate optical access. After surgery the animals were placed in a heated box and monitored to ensure the absence of any sign of discomfort. Before any other experimental procedure, mice were left to recover for 24/48h. During this period, paracetamol (5 mg/ml) was administered in the water as antalgic therapy.

<u>Visual stimulation, data acquisition and analysis</u>: IOS recordings were performed under Isoflurane (1%) and Chlorprothixene (1.25mg/Kg, i.p.). Images were visualized using an Olympus microscope (BX50WI). Red light illumination was provided by 8 red LEDs (625 nm, Knight Lites KSB1385-1P) attached to the objective (Zeiss Plan-NEOFLUAR 5x, NA: 0.16) using a custom-made metal LED holder. The animal was secured under the objective using a ring-shaped neodynium magnet (www.supermagnete.it, R-12-09-1.5-N) mounted on an arduino-based 3D printed imaging chamber that also controls eye shutters and a thermostated heating pad. Visual stimuli were generated using Matlab Psychtoolbox and presented on a gamma corrected 9.7-inch monitor, placed 10 cm away from the eyes of the mouse. Sine wave gratings were presented in the binocular portion of the visual field (−10° to +10° relative to the horizontal midline and −5° to +50° relative to the vertical midline) with a spatial frequency of 0.03 cycles per degree, mean luminance 20 cd/m2 and a contrast of 90%. The stimulus consisted in the abrupt contrast reversal of a grating with a temporal frequency of 4 Hz for 1 sec, time locked with a 16-bit depth acquisition camera (Hamamatsu digital camera C11440) using a parallel port trigger. Interstimulus time was 14 sec. Frames were acquired at 30 fps, with a resolution of 512 x 512 pixels. A total of 270 frames were captured for each trial: 30 before the stimulus as a baseline condition and 240 as post-stimulus. The signal was averaged for at least 30 trials and downsampled to 10 fps. Fluctuations of reflectance (R) for each pixel were computed as the normalized difference from the average baseline (ΔR/R). For each recording, an image representing the mean evoked response was computed by averaging frames between 0.5 to 2.5 sec after stimulation. The mean image was then low-pass filtered with a 2D average spatial filter (30 pixels, 117 μm2 square kernel). To select the binocular portion of the primary visual cortex for further analysis, a region of interest (ROI) was automatically calculated on the mean image of the response by selecting the pixels in the lowest 30% ΔR/R of the range between the maximal and minimal intensity pixel 26. To weaken background fluctuations a manually selected polygonal region of reference (ROR) was subtracted. The ROR was placed where no clear response, blood vessel artifact or irregularities of the skull were observed 27. Mean evoked responses were quantitatively estimated as the average intensity inside the ROI.

### Electrophysiology

<u>Primary neuronal cultures</u>: experiments were performed on cortical neurons obtained from 18-day old embryos of both Cdkl5+/y and Cdkl5^−/y^ mice. The S1 cortex was rapidly dissected under sterile conditions, kept in cold HBSS (4°C) with high glucose, and then digested with papain (0,5 mg/ml) dissolved in HBSS plus DNase (0,1 mg/ml). Isolated cells were then plated at the final density of 1200 cells/mm2. The cells were incubated with 1% penicillin/streptomycin, 1% glutamax, 2.5% fetal bovine serum, 2% B-27 supplemented neurobasal medium in a humidified 5% CO2 atmosphere at 37°C. Experiments were performed at DIV 16 - 18.

<u>Patch-clamp recordings</u>: experiments were performed in voltage clamp conditions and whole-cell configuration as in Marcantoni et al., 2010. Patch electrodes, fabricated from thick borosilicate glasses (Hilgenberg, Mansifield, Germany), were pulled to a final resistance of 3-5 MΩ. Patch Clamp recordings were performed in whole cell configuration using a Multiclamp 700-B amplifier connected to a Digidata 1440 and governed by the pClamp10 software (Axon Instruments, Molecular Devices Ltd, USA). NMDAR activated currents were recorded by holding neurons at −70 mV and perfusing them with the NMDAR agonist, N-Methyl-D-aspartate, (NMDA, 50 μM). The external solution contained (in mM): 130 NaCl, 1.8 CaCl2, 10 HEPES, 10 glucose, 1.2 Glycine (pH 7.4). The internal solution contained (in mM): 90 CsCl, 20 TEACl, 10 glucose, 1 MgCl, 4 ATP, 0,5 GTP, 15 phosphocreatine (pH 7.4). These experiments were performed in the presence of the AMPA and GABAa receptors blockers 6,7-dinitroquinoxaline-2,3-dione, DNQX (20 μM, Sigma-Aldrich) and picrotoxin (100 μM), respectively. Tetrodotoxin (TTX 0.3 μM) was added to block voltage-gated Na+ channels. The mGluR5 were selectively activated for 2 min either by the agonist DHPG (100 μM) or the positive allosteric modulator CDPPB (10 μM).

<u>Miniature post-synaptic currents (mPSCs)</u>: were recorded by holding neurons at −70 mV, recording for 120 seconds, and superfusing the postsynaptic neuron with a Tyrode’s solution containing (in mM): 2 CaCl2, 130 NaCl, 2 MgCl2, 10 HEPES, 4 KCl, and 10 glucose, pH 7.4. The standard internal solution was (in mM): 90 CsCl, 20 TEA-Cl, 10 EGTA, 10 glucose, 1 MgCl2, 4 ATP, 0.5 GTP, and 15 phosphocreatine, pH 7.4. Picrotoxin is added to the Tyrode solution to block GABA A-dependent currents. Tetrodotoxin (TTX, 0.3 μM) will be added for the measure of miniature postsynaptic currents in order to block spontaneous action potentials propagation. Cells were then treated for 2 minutes with DHPG (100 μM). Analysis of peak amplitudes and inter-event intervals (IEI) was performed with Clampfit software (Axon Instruments).

### Behavioural analyses

For the acute treatment regime, 1h after a single i.p. injection with either CDPPB or vehicle, animals were probed with adhesive tape removal test and Y-maze test. For the subchronically treated mice, 24h after the last injection hind-limb clasping behaviour was tested.

<u>Adhesive tape removal test</u>: sensorimotor abilities were evaluated using the adhesive tape removal test as previously described. Briefly, P56 mice were habituated to the testing room for 30 min before starting the experiment and then single animals were placed in the testing cage for the habituation period of 60 sec. The animal was then removed from the testing box and an adhesive tape strip (0.3 cm x 0.4 cm) was placed on the bottom of one forepaw while the other one was lightly touched by the operator with the same pressure. Animals were put back in the testing cage and the latency to touch the tape was recorded with a cut off time of 2 min.

<u>Y-maze test</u>: spontaneous alternation test was used to evaluate spatial working memory in mice. We used an in-house fabricated Y maze that is composed of three arms (34 cm × 5 cm × 10 cm) angled at 120° from one another and made by gray opaque plastic material. Each mouse was placed at the centre of the maze where it can freely explore the three arms for 8 min. Arm entries were defined by the presence of all four paws in an arm. The percentage of spontaneous alternations was calculated as follows: (total alternations / total arm entries − 2) × 100.

<u>Hind-limb clasping</u>: the presence of hind-limb clasping behaviour was tested by suspending the mice from their tail for 2 min and video recorded. Hind-limb clasping scores were assessed as in (Amendola et al., 2014).

### Statistical analysis

All data are reported as mean ± SEM. For the animal experiments, n = number of mice. All statistical analyses were performed using Prism software (Graphpad, La Jolla, CA, USA). For co-IP experiments, one-way analysis of variance (ANOVA) followed by Fisher’s post hoc test was used. For the behavioural and anatomical analyses, Student’s t-test or two-way ANOVA followed by Fisher’s LSD post hoc test were performed, as indicated in the text. For electrophysiology analyses, Student’s t-test and chi-square were used, as indicated in the text. All the raw data are reported in table S1. The statistical analysis performed and the n for each experimental group are reported in figure legends.

## Author’s contribution

AG and MG conceived and designed the study. AG performed the experiments. LL, GS, RM, EP performed IOS experiments. SG performed experiments on human tissues, AG, RP, NM, FP performed behavioural experiments, AG, RP performed immunofluorescence experiments. AM and GC performed electrophysiological experiments. CS, AN synthesized and provided RO6807794; AG, RP, AR, TP, AM and MG analyzed the data. AG and MG wrote the manuscript.

## Institutional Review Board Statement

The study was conducted in accordance with European Community Council Directive 2010/63/UE for care and use of experimental animals with protocols approved by the Italian Minister for Scientific Research (Authorization number 38/2020-PR) and the Bioethics Committee of the University of Torino, Italy.

