## Supplementary material for "mGluR5 PAMs rescue cortical and behavioural defects in a mouse model of CDKL5 deficiency disorder"

[1]Rita Levi-Montalcini" Department of Neuroscience, University of Turin, Turin (Italy); [2]BIO@SNS lab, Scuola Normale Superiore, 56124 Pisa, Italy; [3]Department of Developmental Neuroscience, IRCCS Stella Maris Foundation, 56128 Pisa, Italy, [4]Institute of Neuroscience, CNR, 56124 Pisa, Italy; [5]NEUROFARBA Department, Section of Pharmaceutical and Nutraceutical Sciences, University of Florence, 50019 Sesto Fiorentino (Florence), Italy. [6]Department of Drug Science, University of Turin, Turin, Italy.

Figure S1

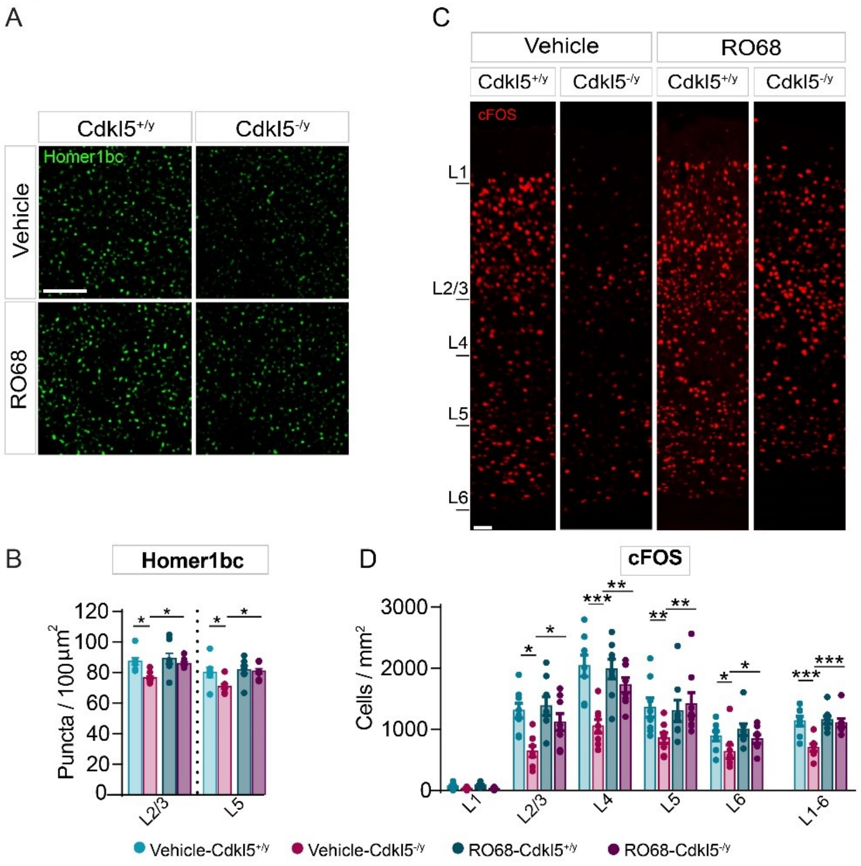


**Figure S1. Acute administration of RO6807794 (RO68) rescues the morpho-functional defects shown by Cdkl5$^{-/y}$ mice.** (**A**) Representative confocal images showing Homer1bc$^{+}$ puncta in layer II-III of S1 cortex from either vehicle- or RO68-treated Cdkl5$^{+/y}$ and Cdkl5$^{-/y}$ mice (scale bar: 5 µm). (**B**) Bar graphs showing differences between genotypes in Homer1bc$^{+}$ immunopuncta density counted from layers II-III and V of S1 cortices in either vehicle- or RO68-treated mice. (**C**) Confocal images of c-FOS$^{+}$ immunostaining in sections of the S1 cortex from Cdkl5$^{+/y}$ and Cdkl5$^{-/y}$ mice, treated with vehicle or RO68 (scale bar: 50 µm), and (**D**) relative cFOS$^{+}$ cells density quantitation throughout layers of V1 cortex. Two-way ANOVA followed by Fisher's multiple comparison test, *$p < 0.05$, ** $p < 0.01$, *** $p < 0.001$; (n = 8 animals for each genotype).

### MATERIALS AND METHODS FOR SUPPLEMENTARY DATA:

#### Chemistry

Solvents and reagents were purchased from Merck and Fluorochem. All reactions involving air- or moisture-sensitive compounds were performed under a nitrogen atmosphere using dried glassware and syringes techniques to transfer solutions. Nuclear magnetic resonance ($^{1}$H-NMR, $^{13}$C-NMR, $^{19}$F-NMR) spectra were recorded using a Bruker Advance III 400 MHz spectrometer in DMSO-$d_6$. Chemical shifts are reported in parts per million (ppm) and the coupling constants (*J*) are expressed in Hertz (Hz). Splitting patterns are designated as follows: s, singlet; d, doublet; t, triplet; m, multiplet; bs, broad singlet. The assignment of exchangeable protons was confirmed by the addition of $D_2O$. Analytical thin-layer chromatography (TLC) was carried out on Merck silica gel F-254 plates. Flash chromatography purifications were performed on Merck silica gel 60 (230-400 mesh ASTM) as the stationary phase and ethyl acetate/n-hexane was used as eluent. The high-resolution mass spectrometry (HRMS) analysis was performed with a Thermo Finnigan LTQ Orbitrap mass spectrometer equipped with an electrospray ionization source (ESI). The analysis was carried out introducing, via syringe pump at 10 μL min$^{-1}$, the sample solution (1.0 μg mL$^{-1}$ in mQ water/acetonitrile 50/50) acquiring the signal of the positive ions. These experimental conditions allowed the monitoring of protonated molecule of the studied compound ($[M+H]^+$ species), that was measured with a proper dwell time to achieve 60 000 units of resolution at Full Width at Half Maximum (FWHM). Elemental compositions of compounds were calculated on the basis of their measured accurate masses, accepting only results with an attribution error less than 2.5 ppm and a not integer RDB (double bond/ring equivalents) value in order to consider only the protonated species.

**Synthesis of N-tert-butyl-5-((3-fluorophenyl)ethynyl)-N-methylpyrimidine-2-carboxamide (RO6807794)**

*Step 1: 5-Bromo-N-(tert-butyl)-N-methylpyrimidine-2-carboxamide.* Oxalyl chloride (1.2 eq) was added dropwise to a suspension of 5-bromopyrimidine-2-carboxylic acid (1 g, 1.0 equiv.) and N,N-dimethylformamide (20 µL) in anhydrous dichloromethane (10 mL). The reaction mixture was stirred for 16h at rt, concentrated in *vacuo* and the obtained residue was dissolved in anhydrous N,N-dimethylformamide (2 mL). Pyridine (1.2 eq) and tert-butyl amine (1.2 eq) were added dropwise to the mixture cooled to 0-5°C and the latter was stirred for 4h at rt. The reaction mixture was quenched with $H_2O$ (15 mL), extracted with ethyl acetate (2 x 30 mL) and the combined organic layers were washed with brine (3 x 20 mL), dried over $Na_2SO_4$, filtered-off and concentrated under *vacuo* to give the pure title compound as a white powder.

*Step 2: N-(Tert-butyl)-5-iodo-N-methylpyrimidine-2-carboxamide.* Sodium iodide (4 eq), copper(I) iodide (0.2 eq) and trans-N,N'-dimethylcyclohexane-1,2-diamine (0.2 eq) were added to a solution of 5-bromo-N-(tert-butyl)-N-methylpyrimidine-2-carboxamide (from *Step 1*, 0.8 g, 1.0 eq) in dioxane (10 mL). The reaction mixture was stirred for 16h at 100°C until consumption of the starting materials (TLC monitoring), cooled to rt, quenched with a saturated $NaHCO_3$ solution (30 mL) and extracted with ethyl acetate (3 x 30 mL). The combined organic layers were washed with brine (3 x 25 mL), dried over $Na_2SO_4$, filtered-off and concentrated under *vacuo* to give a residue that was purified by silica gel column chromatography eluting with ethyl acetate/n-hexane 20 to 50 % yielding the pure title compound as a white powder.

*Step 3: N-Tert-butyl-5-((3-fluorophenyl)ethynyl)-N-methylpyrimidine-2-carboxamide.* Bis-(triphenylphosphine)palladium (II) dichloride (0.1 eq), triethylamine (3.0 eq), triphenylphosphine (0.1 eq) and copper(I) iodide (0.1 eq) were added to a solution of N-(tert-butyl)-5-iodo-N-methylpyrimidine-2-carboxamide (from *Step 2*, 0.6 g, 1.0 eq) and 3-fluorophenylacetylene (2.0 eq) in anhydrous N,N-dimethylformamide (2 mL). The reaction mixture was stirred for 2h at 70°C, cooled to rt, quenched with a saturated $NaHCO_3$ solution (15 mL) and extracted with dichloromethane (3 x 20 mL). The combined organic layers were washed with brine (3 x 15 mL), dried over $Na_2SO_4$, filtered-off and concentrated under *vacuo* to give the raw derivative that was purified by silica gel column chromatography eluting with ethyl acetate/n-hexane 20 to 40 % affording the pure title compound (RO680794) as a light grey powder: 58% yield; TLC $R_f$ 0.30 (ethyl acetate/n-hexane 40 % v/v); $^1$H-NMR (DMSO-d$_6$, 400 MHz): δ 1.47 (9H, s, 3 x C$H_3$), 2.67 (3H, s, C$H_3$), 7.37 (1H, m, Ar-*H*), 7.53 (3H, m, Ar-*H*), 9.07 (2H, s, Ar-*H*); $^{19}$F-NMR (376 MHz, DMSO-d$_6$): -112.09 (s, 1F); $^{13}$C-NMR (DMSO-d$_6$, 100 MHz): δ 28.19, 34.12, 57.37, 84.63, 95.29, 118.04 (d, $J^2_{CF}$ = 21.0), 118.41, 119.16 (d, $J^2_{CF}$ = 24.0), 123.97 (d, $J^3_{CF}$ = 10.0), 128.96, 132.09 (d, $J^3_{CF}$ = 9.0), 160.30, 162.41, 162.77 (d, $J^1_{CF}$ = 243.0), 167.44; ESI-HRMS (m/z) $[M+H]^+$: calculated for C18H19FN3O 312.1512; found 312.1508 (Jaeschke et al. 2012)

**Drug administration:** 0.3 mg/kg of RO6807794 was injected intraperitonially, as reported by Kelly et al., 2018. The drug was prepared always fresh by dissolving it in 0.3% of Tween 80 and the volume was made up with normal saline (Kelly et al., 2018). The experimental plan included four groups of mice, i.e.: WT mice treated with vehicle; WT mice treated with RO68; Cdkl5 KO mice treated with vehicle and Cdkl5 KO mice treated with RO68. Two hours after the treatment, the animals were sacrificed, and brains collected for immunofluorescence.

**Immunofluorescence and Images acquisition and analysis**: see the relevant methods section in the main text.
